# Beyond Animals and Plants: Dynamic Maternal Effects in the Fungus *Neurospora crassa*

**DOI:** 10.1101/028092

**Authors:** Kolea C. K. Zimmerman, Daniel A. Levitis, Anne Pringle

## Abstract

Maternal effects are widely documented in animals and plants, but not in fungi or other eukaryotes. A principal cause of maternal effects is asymmetrical parental investment in a zygote, creating greater maternal versus paternal influence on offspring phenotypes. Asymmetrical investments are not limited to animals and plants, but are also prevalent in fungi and groups including apicocomplexans, dinoflagellates, and red algae. Evidence suggesting maternal effects among fungi is sparse and anecdotal. In an experiment designed to test for maternal effects across sexual reproduction in the model fungus *Neurospora crassa*, we measured offspring phenotypes from crosses of all possible pairs of 22 individuals. Crosses encompassed reciprocals of 11 mating-type “A” and 11 mating-type “a” wild strains. After controlling for the genetic and geographic distances between strains in any individual cross, we found strong evidence for maternal control of sporocarp production, as well as maternal effects on spore numbers and spore germination. However, both parents exert equal influence on the percentage of spores that are pigmented, and size of pigmented spores. We propose a model linking the stage-specific presence or absence of maternal effects to cellular developmental processes: effects appear to be mediated primarily through the maternal cytoplasm, and, after spore cell walls form, maternal influence on spore development is limited. Maternal effects in fungi, thus far largely ignored, are likely to shape species’ evolution and ecology. The association of anisogamy and maternal effects in a fungus suggests maternal effects may also influence the biology of other anisogamous eukaryotes.

## INTRODUCTION

Mothers influence offspring phenotypes not only by transmitting genes but also by directly altering the course of offspring development. Maternal influences extend beyond inherited nuclear chromosomes. For example, they include cytoplasmic provisioning and the environment provided by the mother during embryogenesis. These are named maternal effects (Roach & Wulff, 1987). The consequences of maternal effects for evolution are diverse. Maternal effects can alter the direction and strength of trait selection by introducing sources of variation outside of an individual’s genes (Kirkpatrick & Lande, 1989). Maternal effects enable the environmental influences on mothers to alter the development of her offspring, effectively transferring environmental influences across generations (Galloway, 2005; Badyaev & Uller, 2009; Donohue, 2009). In clonal organisms, environmental maternal effects can increase offspring heterogeneity (Sakwinska, 2004).

A basis for maternal effects can be traced to a common phenomenon apparent in most animals and plants: the female gamete is much larger than the male gamete, enabling a greater female influence on offspring (Westneat & Craig Sargent, 1996). When differently sized gametes fuse, maternally derived cytoplasm predominates in the zygote (maternal cytoplasmic inheritance) and disproportionately influences zygote development (Birky, 2001; Gosden, 2002). Maternal effects are also found in species where offspring develop inside or on the mother, for example, where an animal develops inside a womb (Lindström, 1999), or seeds develop in a fruit attached to a plant (Roach & Wulff, 1987).

While breeders have long known that mothers exert more influence on offspring than fathers (Brooks, 1905), maternal effects were first carefully described using quantitative approaches in animals (Walton & Hammond, 1938) and more recently in plants (Roach & Wulff, 1987; Mousseau *et al.*, 2009). Maternal effects are rarely discussed outside of these kingdoms. The basic mechanisms causing maternal effects in species of these paradigmatic kingdoms— anisogamy leading to disproportionate cytoplasmic inheritance and offspring development in maternally derived environments—are also found among a variety of other eukaryotes, including apicocomplexans, dinoflagellates, diatoms, red algae (Dacks & Kasinsky, 1999), and fungi (Jinks, 1963; Billiard *et al.*, 2010; Debuchy *et al.*, 2010). Within the fungi, the prerequisites for maternal effects are commonly found among filamentous species in the Ascomycota, and more rarely, in the Basidiomycota (Billiard *et al.*, 2010). Most species in the Ascomycota have distinct male and female structures, whereas there is very little morphological specialization of sex organs in the Basidiomycota. Anisogamy involving unequal parental investment in offspring has been proposed as a basis for sexual selection in fungi (Nieuwenhuis & Aanen, 2012).

The genetic model *Neurospora crassa* (Fig. 1) is a well characterized and experimentally tractable filamentous Ascomycete (Roche *et al.*, 2014), and it provides an optimal system for tests of maternal effects in fungi. The female gamete-forming organs (protoperithecia) are much larger (up to 100 μm in diameter) than the male gametes (2.5-8.5 p,m in diameter) and offspring development takes place in a maternally produced “fruiting body,” or sporocarp (perithecium). *N. crassa* is hermaphroditic, enabling reciprocal crosses to test for maternal effects.

**Figure 1.**
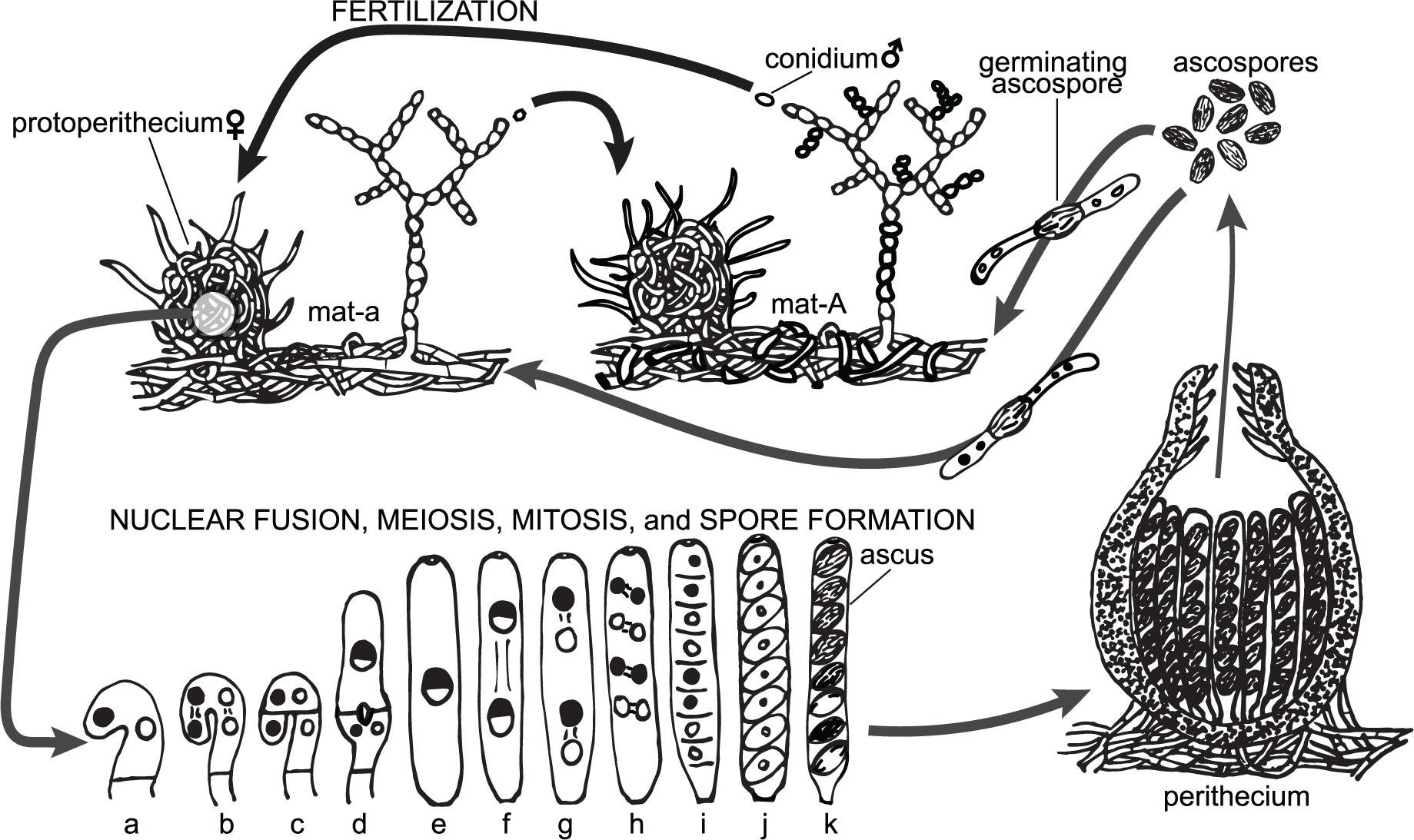
Life cycle of *Neurospora crassa.* *N. crassa* is a haploid filamentous fungus capable of reproducing both sexually and asexually (Davis, 2000). Sexual reproduction is restricted to parent strains different at a single two-allele mating-type locus and a single strain is either mat-a or mat-A. Both mating-types can produce female and male gametes; all strains are hermaphroditic. For example, reciprocal fertilization between mat-a and mat-A strains is shown at the top of the figure. Nitrogen starvation induces the production of protoperithecia, which are female reproductive structures that house female gametes termed ascogonia. Asexual spores termed conidia (or potentially other cells from a compatible mate) act as male gametes. Fertilization follows the growth of a trichogyne—a sexually receptive protoperithecial hypha— and its fusion with a conidium of the opposite mating type. After fusion, the paternal nucleus migrates to an ascogonium within the protoperithecium. The division and segregation of maternal and paternal nuclei results in the proliferation of cells each containing one maternal and one paternal nucleus (**a-c**). Within these cells, each of the two nuclei fuse to form a diploid zygote (**d, e**) that rapidly undergoes meiosis I (**f**) and meiosis II (**g**), producing four haploid nuclei. These nuclei then undergo mitosis, resulting in eight haploid nuclei per ascus (**h**). Cytoplasmic cleavage occurs (**i**) and a cell wall develops around each of the eight haploid nuclei, forming eight ascospores (**j**). In normal ascospores, the cell wall becomes pigmented as it matures; in aborted or undeveloped ascospores, the cell wall remains unpigmented (**k**). Spores are forcibly ejected from a mature perithecium one ascus at a time. Nuclei are colored black or white to show that they are derived from the mat-a or mat-A parent, respectively. All nuclei are haploid (1N) except for the diploid (2N) half-black/half-white nuclei shown in steps **d, e**, and **f**.

To our knowledge, no breeding or quantitative genetics experiments have investigated maternal effects in *N. crassa* or any other fungus, but *post hoc* evidence of reciprocal differences in fertility among *N. crassa* strains suggests the presence of strong maternal control over reproductive development. While every strain of *N. crassa* is hermaphroditic—and can function as either a female or a male or both—there are mutant *N. crassa* strains with phenotypes that are only apparent when the strain acts as the maternal parent; these kinds of mutations cause defects in female gamete formation or fertilization and result in reciprocal differences in fertility (Raju, 1992). Experiments with wild strains of *N. crassa* also document reciprocal differences in fitness, specifically, the ability to form perithecia suggests protoperithecial formation and fertilization are influenced by the maternal parent even among wild fungi (Dettman *et al.*, 2003b; Turner *et al.*, 2010). Reciprocal crosses between experimentally evolved *Neurospora* strains also show maternal influence on mating success (Dettman *et al.*, 2008).

Studies to date prove that female structures (protoperithecia and perithecia) in *N. crassa* are truly female and play a dominant role in the sexual cycle of this fungus. However, by definition, maternal effects act on offspring and can only be determined if offspring and their traits are measured. Although many mutations affecting the production of sexual spores (termed ascospores, and hereafter referred to as “spores”) have been identified in *N. crassa*, these mutations either segregate after meiosis or are only expressed in crosses homozygous for the allele, indicating they are direct genetic effects and not maternal effects (Leslie & Raju, 1985; Raju *et al.*, 1987; Raju & Leslie, 1992).

Experiments testing for maternal effects among fungi are long overdue. Fungi are critical drivers of ecosystem function, with roles as pathogens, mutualists, and decomposers. The diverse evolutionary consequences of maternal effects documented among animals and plants are likely equally relevant to anisogamous fungi. The majority of fungal pathogens are Ascomycetes (Guarro *et al.*, 1999) and many of the most damaging plant diseases, including rice blast *(Magnaporthe oryzae)* and grey mould *(Botrytis cinerea)* (Dean *et al.*, 2012) have life cycles very similar to the life cycle of *N. crassa* (Faretra *et al.*, 1988; Saleh *et al.*, 2012). If maternal effects play a role in the evolutionary trajectories or ecological niches of these economically relevant species, documenting the phenomena involved may provide novel tools to understand pathogen biology. Moreover, testing for maternal effects among fungi provides an opportunity to extend theory generated from two groups, animals and plants, to diverse eukaryotes.

To test whether maternal effects occur in *Neurospora crassa*, we mated genotyped strains in a fully and reciprocally crossed experiment. Strains were originally isolated from a diversity of sites across the globe. We used generalized linear mixed effects models to quantify the effects of crossing distances (the genetic and geographic distances between parents in any particular cross) and the variation in strains’ influence, as mothers or fathers, on traits (Table 1) associated with the sexual ontogeny of the fungus. Geographic and genetic differences among strains may influence reproductive success because, for example, crossing closely related parents results in inbreeding depression or crossing distantly related parents results in outbreeding depression (Lynch, 1991). Our analyses document clear maternal effects on offspring phenotypes, but the presence and strength of these effects varies dynamically across development.

**Table 1.** Target traits.

| **Trait** | **Description** | **Characteristic of:** |
| --- | --- | --- |
| Perithecia Count | Number of protoperithecia fertilized: perithecia are sporocarps housing zygotes; zygotes immediately undergo meiosis to form spores. | Mother |
| Spore Count | The spores produced by a given cross; each zygote has the potential to result in 8 spores. | Zygotes of a particular cross |
| Proportion Pigmented | Proportion of spores that are pigmented. Analogous to measuring the potential gametic viability of the F1 generation; pigmented spores are mature and potentially viable while all unpigmented spores are inviable. | Spores of a particular cross |
| Pigmented Spore Size | Mean length of pigmented spores. | Spores of a particular cross |
| Pigmented Spore Germination | Proportion of pigmented spores that germinate within 150 mins. after activation of spores. | Spores of a particular cross |

## METHODS

### Mating scheme and strain choice

We used a full diallel cross design to quantify parental effects in *N. crassa.* In a full diallel, all possible combinations of parents are mated, along with reciprocals *(i.e.* each strain is used as a mother, then a father, with every other compatible strain). The mating design maximizes the statistical power for estimating maternal, paternal, and direct genetic effects on offspring (Zhu & Weir, 1996). To control for the effects of crossing distances on reproductive output, we first identified a set of 24 mat-A and 24 mat-a wild isolates of *N. crassa* (Fig. 2, Table 2), previously genotyped with RNAseq (Ellison *et al.*, 2011).

**Figure 2.**
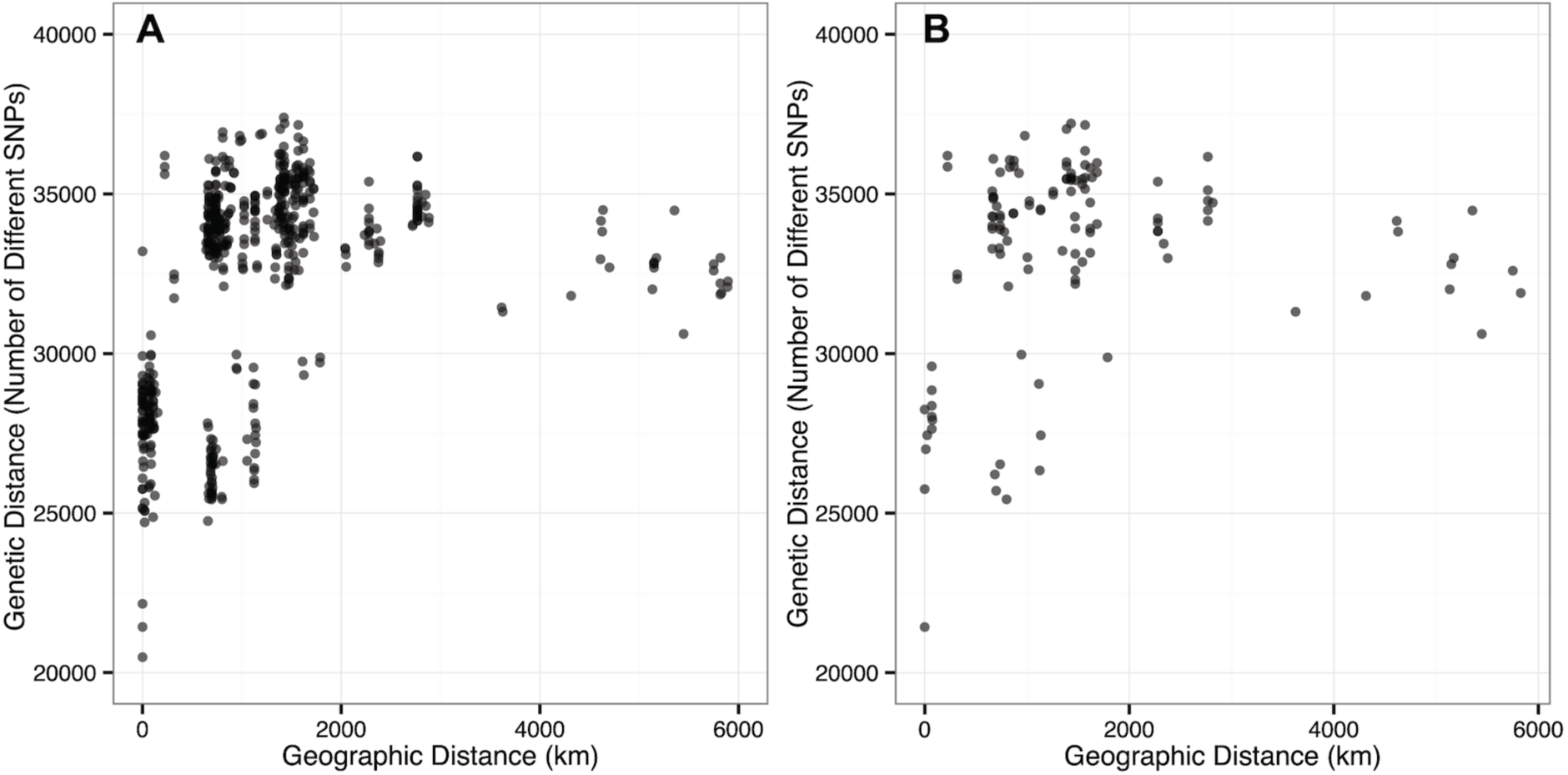
Experimental crosses chosen from among all potential crosses using the SPREAD algorithm. (A) Genetic distance *vs.* geographic distance between parents of all possible crosses. (B) the subset of 11 mat-A x 11 mat-a crosses selected with the SPREAD algorithm (Zimmerman *et al.*, 2015). Crosses are plotted as semitransparent dots and darker shading marks overlapping crosses.

**Table 2.**
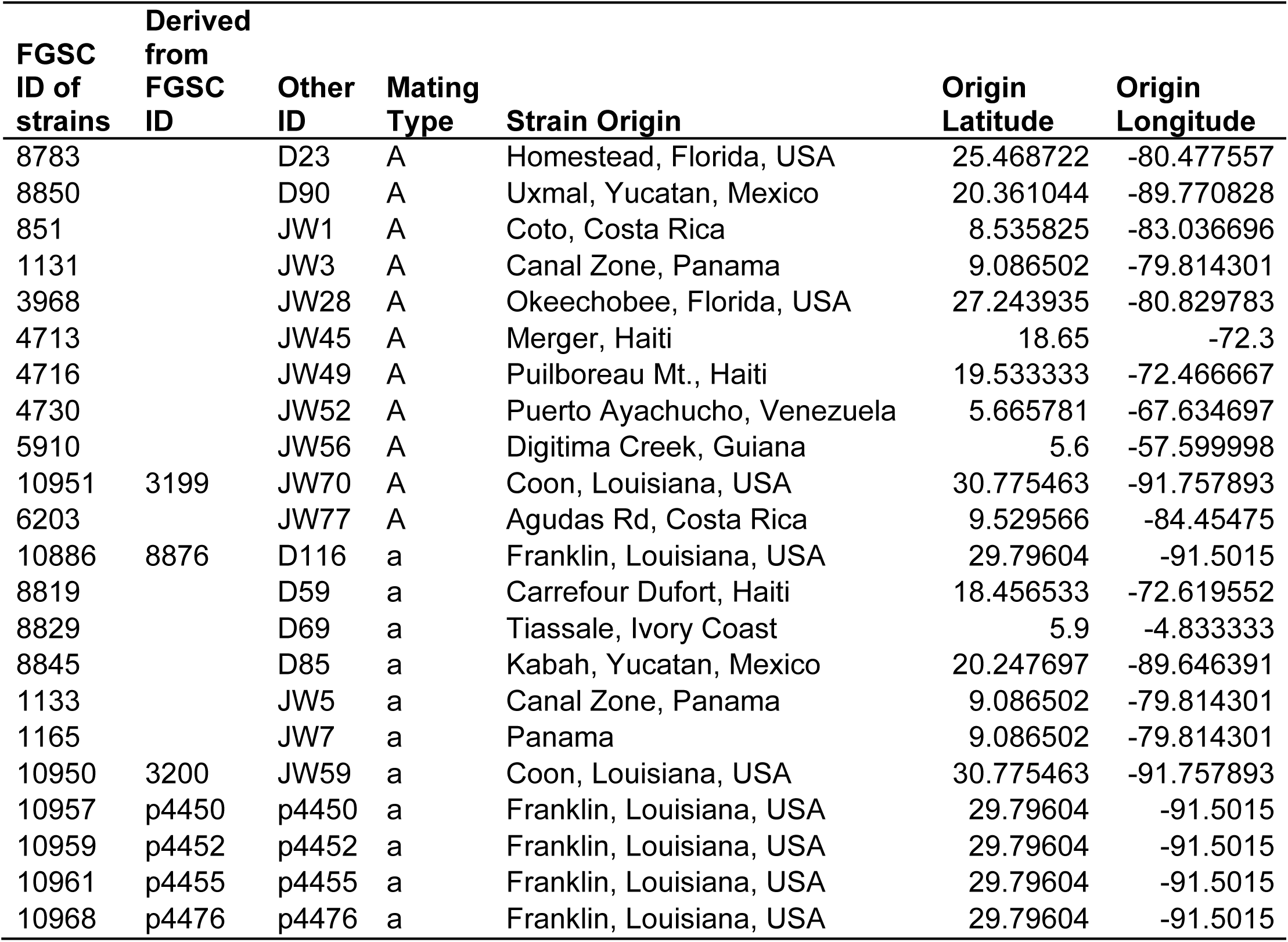
Experimental *N. crassa* strains. Strains used in this experiment are shown in the first column, “Derived from FGSC ID” = ID of heterokaryotic strain purified to homokaryons by Ellison et al. (Ellison *et al.*, 2011), “Other ID” = additional names for the same strain used in other publications.

We used the SNP data (Ellison *et al.*, 2011) to calculate pairwise genetic distances between all possible pairs of strains, measured as number of differences across all SNP sites. For each of the 48 strains, we obtained collection site information from the Fungal Genetic Stocks Center strain catalog (FGSC, www.fgsc.net) and calculated geographic distances between strain collection sites.

The genotyped population included 24 strains of each mating type, but generating all possible crosses with these 48 strains (a total of 576 crosses) was beyond our experimental capacity. Using the SPREAD algorithm (Zimmerman *et al.*, 2015), we selected a mating set of 11 mat-a and 11 mat-A strains for use in our study. SPREAD identifies the subset of crosses most likely to mimic the original distribution of the entire set of all potential crosses. Briefly: SPREAD selects a crossing-set from 1000 randomly generated crossing-sets (11 mat-a × 11 mat-A) by maximizing the mean nearest neighbor distance of the subset of 121 crosses plotted according to the geographic *vs*. genetic distances among crosses. We obtained the 22 strains chosen by this algorithm (Table 2) from the FGSC (McCluskey *et al.*, 2010).

### Generating conidia for crosses

We inoculated each strain onto a slant of Vogel’s Medium N (Vogel, 1956) made with 2% sucrose and 1.5% agar. We harvested conidia from these cultures and preserved them on silica gel for subsequent experiments. To grow conidia for crosses we used conidial stocks to inoculate 50 mL narrow neck flasks containing 7 mL of Vogel’s Medium N with 2% sucrose, 1.5% agar, 0.1% yeast extract, and 0.1% casamino acids, incubated at 30°C with 12h/12h light/dark illumination for 7 days. We harvested conidial inocula for crosses with 10 mL diH_2_O and filtered suspensions through non-absorbent cotton to remove extraneous mycelia.

### Crossing conditions and spore collection

We grew four replicate reciprocal crosses of each of the 121 strain pairs for a total of (4 replicates) × (121 mating pairs) × (2 reciprocals) = 968 crosses. We performed all crosses using paper crossing media (Pandit & Russo, 1993) with slight modifications: we prepared crossing plates by placing 4 × 3.5cm gel blot paper squares in 5 cm petri dishes and adding 1 mL of synthetic crossing media (Westergaard & Mitchell, 1947) containing 0.1% sucrose, 0.5% sorbose, 0.1% yeast extract and 0.1% casamino acids. We grew each of the 22 strains as a female by evenly dropping 200 μL of a suspension containing 1 × 10^6^ conidia in diH_2_O onto the paper. We incubated plates in their plastic sleeves in the dark, at 25°C for 48 hours, and then fertilized the plates by evenly dropping 200 μL of a suspension containing 1x10^6^ conidia in diH_2_O onto the filter paper. Because these conidia are dropped onto individuals already growing as females, they will function as the male gametes. We incubated the fertilized crosses at 25°C in darkness for 20 days, except for occasional exposures to light during routine inspections. After incubation, we harvested spores from lids with 1mL of diH_2_O and deposited spores in deep 96-well plates. To allow for complete maturation of the spores, and to standardize spore age among crosses, we stored the harvested spores at 4°C for 16 days before flow cytometry analyses of spore counts, morphology, and germination.

### Counting perithecia

We photographed all crosses at time of ascospore harvest for subsequent counting of perithecia. Using the photographs, we determined total counts of perithecia per cross by manual counting and marking in ImageJ (Schneider *et al.*, 2012).

### Counting and measuring spores using flow cytometry

Due to the large number of spore samples to be analyzed, we used high-throughput flow cytometry to characterize spore samples collected from the crosses. In a single run of a spore sample through the flow cytometer, we counted spores in the sample and measured the size and pigmentation of spores. Using a high-throughput sampling accessory, we were able to analyze spore samples at a rate of 1 sample/minute.

We prepared spores for flow cytometry by first centrifuging the 96-well plates with harvested spore suspensions at 500 rpm for 10 mins. We pipetted away the supernatant and resuspended ascospores in 500 μL of 1% wt/v polyvinylpyrrolidone molecular mass 360,000 (PVP-360) in diH_2_O. We transferred 50 μL of this ascospore suspension, along with 100 μL of diH_2_O to new 96-well U-bottom plates. To wash PVP-360 out of suspensions, we centrifuged the plates for 5 mins at 1000rpm, then removed 125 μL of the supernatant and added 125 μL diH_2_O. Next, we added Spherotech Accucount 7.7 μm fluorescent beads (Spherotech Inc., Lake Forest, IL) to the spore suspension in 50 μL of 0.4% Triton X-100 (Sigma Chemical Co., St. Louis, MO), resulting in a final spore and bead suspension in 200 μL of 0.1% Triton X-100 in diH_2_O. Spore samples were analyzed by a BD LSRII flow cytometer with a high throughput sorter attachment (BD Biosciences). The sampling parameters for high throughput sorting were as follows: Sample flow rate: 3.0 μL/s, sample volume: 50 μL, mix volume: 100 μL, mix speed: 75 μL/s, number of mixes: 5, wash volume: 300 μL. We collected data on three parameters: forward scatter (to measure pigmentation), side scatter (to measure size), and fluorescence (to measure bead size). Using FACSDiVa software (BD Biosciences), we recorded data on forward scatter height, width, and area; side scatter height, width, and area; and fluorescence height, width, and area for every spore in the sample volume (50 μL). The data from each sample were saved in the Flow Cytometry Standard (FCS) 3.0 format.

### Measuring spore germination with flow cytometry

To measure the relative germination abilities of spores collected from all crosses, we used flow cytometry to measure the length of spores in spore samples from each cross before and after exposure to germination triggers (nutrient media and heat-shock). As spores germinate, spore length increases because germ tubes emerge from each end of the spore. For each sample, we transferred 50 μL of the spore suspension in 1% PVP-360 to duplicate U-bottom 96-well plates, each representing either pre-germination or post-germination treatments. Each well contained 100 μL of diH_2_O and plates were centrifuged for 5 mins at 1000 rpm. We removed 100 μL of supernatant and added 50 μL of 2x Vogel’s Medium N containing 4% sucrose, resulting in a final concentration of 1x Vogel’s Medium N containing 2% sucrose. To control for the effects of heat-shock on spore size (pre-germination plate) and activate germination (post-germination plate), we heat-shocked both plates in a forced air incubator at 60°C for 1 hour. We fixed the pre-germination plates immediately following heat-shock. Post-germination plates were incubated with shaking for 150 mins. at 34°C to promote germ tube growth, and then fixed.

We fixed spores by adding 100 μL of cold 95% EtOH and 0.2% Triton X-100 in diH_2_O directly to the spore suspensions, and incubating overnight at 4°C. Just before flow cytometry, we transferred spores to 96-well PCR plates and centrifuged them at 2000 rpm for 10 mins. We removed 175 μL of the supernatant and added 175 μL of diH_2_O, resuspended spores by gentle pipetting, and transferred the resuspended spores to new 96-well U-bottom plates along with 25 μL of a Spherotech 7.7 μm fluorescent bead solution containing ~ 2550 beads in 0.4% Triton X-100. Flow cytometry parameters were the same as above.

### Flow cytometry data processing

Our initial analysis of the flow cytometry data indicated the presence of both conidia and ascospores, with ascospores exhibiting two clear sub-populations of pigmented and hyaline (unpigmented) ascospores (Fig. S1). Hyaline ascospores are demarcated by greater forward scatter height values, compared to pigmented ascospores; because they are clear they scatter light differently and exhibit higher light transmittance. There was some overlap of conidia and ascospores when plotted on forward-scatter-height *vs.* side-scatter-width axes, and the overlap prevented manual two-dimensional gating. In order to differentiate the conidia and ascospores, we performed multi-dimensional clustering analyses with all measured flow cytometery parameters, using a custom clustering and sorting pipeline in R with the following packages: *flowCore* (Ellis *et al.*, n.d.), *flowClust* (Lo *et al.*, 2008; 2009), and *flowStats* (Hahne *et al.*, n.d.).

FCS files were imported into R using *flowCore* and sets of FCS files for each plate were merged into a single dataset to facilitate clustering by increasing the density of measurements. To distinguish the populations of conidia, pigmented ascospores, and hyaline ascospores, we performed k-means clustering using the *flowClust* package. We manually adjusted the number of clusters to be identified for each dataset to obtain clustered datasets whose clusters matched sub-population characteristics obtained from analyzing pure conidia or pure ascospores. We then computed summary statistics using *flowStats* for each measured sample including the total number of ascospores, the number of pigmented spores, the number of hyaline ascospores, and the size of pigmented ascospores. Fluorescent bead populations were identified based on their high fluorescence parameter values (fluorescence-area). To standardize ascospore counts in subsequent analyses, we used the flow cytometry sample measurement time of each sample. To standardize spore size across samples we used the mean side-scatter width of the fluorescent beads for each sample.

To calculate ascospore germination, we first drew rectangular gates around populations of spores from pre-germination ascospore samples plotted on forward scatter height *vs*. side scatter width. We applied these gates, representing the size of pre-germination ascospores, to the post-germination ascospore samples and counted the ascospores outside of the previously drawn gates as germinated ascospores. For each pre-germination sample, we used our clustering pipeline to determine the proportion of conidia and ascospores, excluding fluorescent beads. Using these proportions, we calculated the total ascospore number in each post-germination sample (including germinated ascospores) by multiplying the proportion of total ascospores in the pre-germination sample by the total number of particles (excluding fluorescent beads) in the post-germination sample. Finally, we calculated the proportion of germinated ascospores by dividing the number of germinated ascospores (determined by gating) by the estimated total number of ascospores in the sample (determined by the clustering analysis of the pregermination sample).

### Verification of flow cytometry methods

We verified the ability of our flow cytometry analysis to measure ascospore pigmentation and germination by performing traditional counting and plating assays for a subsample of the crosses used in the full crossing experiment. In a separate experiment, we replicated a subset of crosses and measured ascospore production, pigmentation and germination using flow cytometry as described above. Then, using the same ascospore samples, we obtained counts of pigmented and hyaline ascospores by dropping 10 μL of a spore suspension onto plates (Vogel’s medium N with 1% sucrose and 1.5% agar) and examining the drop under a dissecting microscope. Next, we spread the spores using a bent Pasteur pipette (to minimize spore loss), heatshocked the plates for 1 hr. at 60°C, and incubated them overnight at 30°C. To measure the number of germinated spores, we counted growing colonies using a dissecting microscope. We determined the strength and nature of the relationship between measurements obtained from flow cytometry and manual methods using asymptotic non-linear least squares regression, with the SSasymp function from the *stats* package in R. Analyses confirm a strong relationship between the two different methods of measuring spores, confirming that flow cytometry can be used as a proxy for manual counting (Figs. S2 and S3).

### Mixed effects models

We used the R package *glmmADMB* to compute mixed effects models for each trait (Fournier *et al.*, 2012; Skaug *et al.*, 2013). Based on preliminary analyses of the trait data distributions, we chose to model pigmented spore size with a Gaussian distribution and all other response variables with a negative binomial distribution. Our initial models included the following fixed effects: genetic distance (number of different SNPs between parents in a cross), geographic distance (km between strain locales), the squares of both genetic and geographic distance, and an interaction term for genetic and geographic distance. In each model, we also included maternal and paternal parentage as random effects, specifically to test the hypothesis that strains will vary in their ability, as mothers or fathers, to influence the target traits. For all five phenotypic traits we computed full models and then selected simplified final models using the quasi Akaike Information Criterion (qAIC) or Akaike Information Criterion (AIC) for models with negative binomial or Gaussian response variable distributions, respectively.

Best linear unbiased predictors (BLUPs), also referred to as random effect coefficients or conditional modes, are model based estimates of the relative influence of a strain as a mother or a father on a given phenotype, after controlling for fixed effects. For each trait, we calculated the relative proportions of overall random effect variance attributed to the variance of maternal or paternal BLUPs and assessed correlations between maternal and paternal BLUPs.

### R^2^ GLMM calculations

To determine the relative importance of crossing distances (fixed effects) and the identity and sex of the parents (random effects) in selected models, we calculated the proportion of variance explained by either fixed effects or just random effects for each model using R^2^GLMM statistics (Nakagawa and Schielzeth 2013). For the pigmented spore size model, we used equations for calculating R^2^GLMM statistics for models with a Gaussian response distribution listed by Nakagawa and Schielzeth (2013). For the other models that use negative binomial response distributions, we derived the distribution specific variance 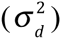 for negative binomial distributions with a log link function where *α* is the negative binomial dispersion parameter following the derivation in Appendix 1 of Nakagawa and Schielzeth (2013). Our derivation is shown in the supplementary information (Derivation S1).

### Highest posterior density of parameter estimates

We verified the significance of parameter values in each selected model by calculating HPD intervals for parameter values. HPD intervals, also known as Bayesian confidence intervals, are based on the observed data and indicate the range of values in which the true parameter value is found with a 95% probability. HPD intervals of model parameter values that do not span zero are deemed significant. We calculated HPD intervals of the parameters in each model using the built-in Markov chain Monte Carlo (MCMC) functionality of glmmADMB that uses parameter estimates calculated from the model as priors. We used a chain of 10000 MCMC iterations and verified MCMC chain convergence by plotting the distributions of samples drawn from the chain. Parameter estimates with HPD intervals that did not span zero were deemed significant.

## RESULTS

### Offspring phenotypes from a full diallel cross show evidence of maternal effects in *N.crassa*

All crosses resulted in perithecia and viable spores, except crosses involving strain FGSC 10968 (mat-a), which was infertile as both male and female and exhibited abnormal colony growth with increased pigmentation of mycelia. We excluded crosses involving FGSC 10968 from all subsequent analyses.

Crosses involving different strains resulted in a diverse array of trait phenotypes, and differences between reciprocal crosses were observed for most traits (Fig. 3). When crosses involving a particular strain acting as a female are grouped, the maternal effects of different strains are obvious. For example, when strain 10951 (mat-A) is used as a female, crosses appear to have the same level of spore production no matter the identity of the male strain (Fig. 3C). These kinds of patterns are also observed for data on pigmented spore germination, see for example crosses involving 8850 (Fig. 3I) or 1165 (Fig. 3J). But when crosses involving a particular strain acting as a male are grouped, patterns are more variable; for example, the influence of strain 8783 (mat-A) when it is used as a male depends on the pairing: with 8845 (mat-a) many spores are produced, but with 1165 (mat-a), few spores are produced (Fig. 3D). These patterns: a consistent maternal influence, and variable paternal influence, are not easily observed within the data for proportion pigmented (Fig. 3E, F) or pigmented spore size (Fig. 3G, H).

**Figure 3.**
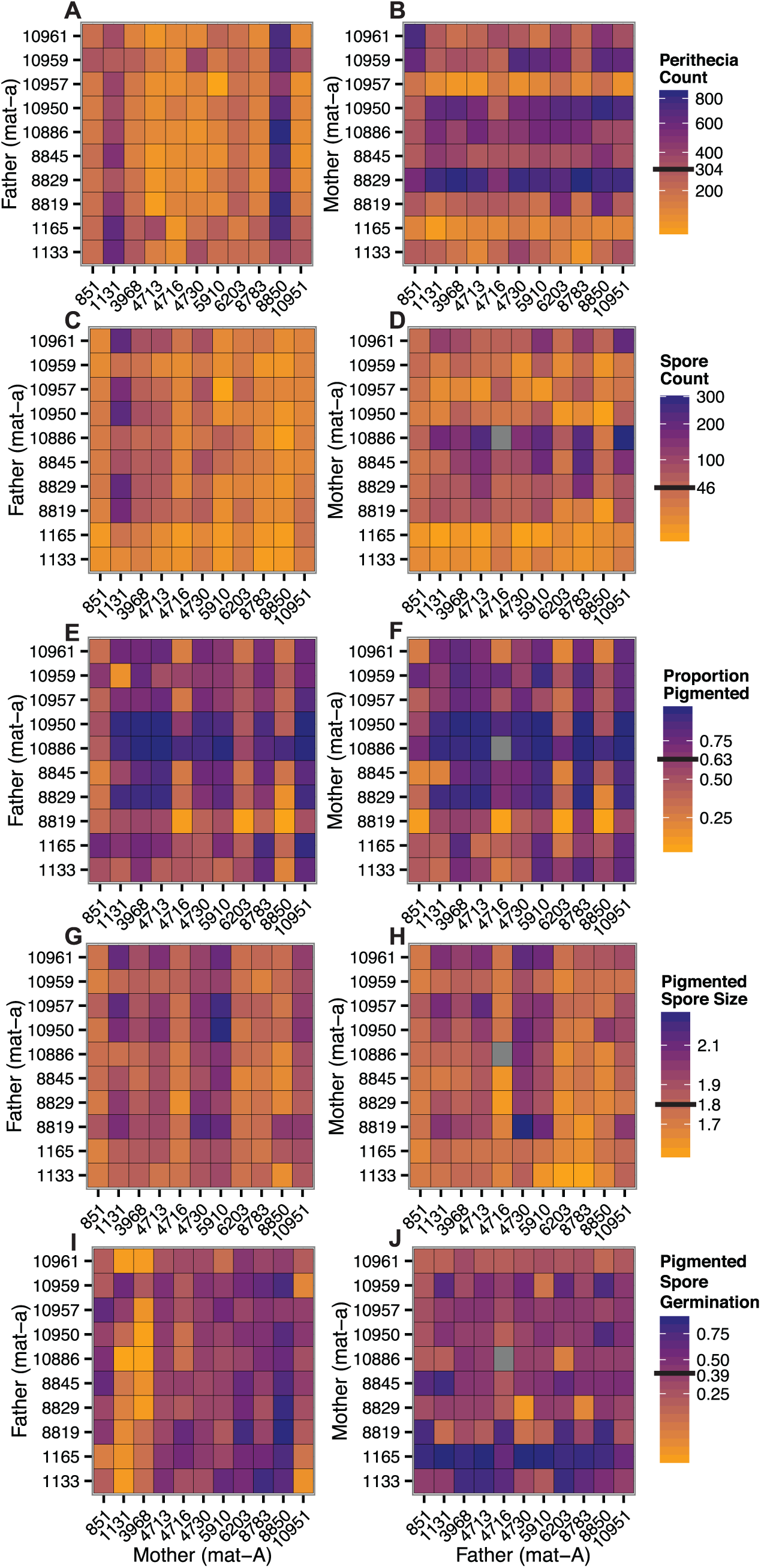
Trait phenotypes for all crosses. Heatmaps show crosses between mat-a fathers and mat-A mothers (panels A, C, E, G, and I) and reciprocal crosses between mat-a mothers and mat-A fathers (panels B, D, F, H, and J). Scales at right are values for a particular trait; black bars on scales are mean values for the trait, calculated from all crosses. Grey boxes mark missing data caused by instrument error.

Whether reciprocal crosses result in different phenotypes depends on the traits involved. Spore count and pigmented spore germination data are often different between reciprocal crosses, for example few spores are produced by crossing 10951 as female and 10961 as male (Fig. 3C), but many spores are produced by crossing 10951 as male with 10961 as female (Fig. 3D). However, there is very little difference in the proportion of pigmented spores (Fig. 3E, F) or pigmented spore size (Fig. 3G, H) between reciprocal crosses, as shown by the almost identical reciprocal heatmaps for these traits.

### Mixed effects models confirm parental effects exert greater influence on trait outcomes than crossing distances

Visual inspection of trait phenotypes suggests that mothers have more control over spore production and spore germination than fathers. But to more carefully explore the relationship between spore characteristics and the multiple influences of parental effects and crossing distances, we constructed generalized linear mixed effects models for each measured trait (Table 3). Highest posterior density intervals for the selected parameters indicate that parameters for all models are significantly different from zero, except the genetic distance parameter in the spore count model (Fig. 4).

**Table 3.** Summary of best models. By default, all models include the random effects of maternal and paternal parentage.

| Trait | Equation | Distribution | Link | Offset | Model Selection Method |
| --- | --- | --- | --- | --- | --- |
| Perithecia Count | Perithecia Count ~ Genetic Distance + (Genetic Distance)<sup>2</sup> | Negative Binomial | log | --- | qAIC |
| Spore Count | Total Spore Count ~ Genetic Distance + (Genetic Distance)<sup>2</sup> + Geographic Distance | Negative Binomial | log | Sample Run time (FCS) | qAIC |
| Proportion Pigmented | Pigmented Spore Count ~ Genetic Distance + Geographic Distance + (Geographic Distance)<sup>2</sup> + (Genetic Distance)<sup>2</sup> + Genetic Distance: Geographic Distance | Negative Binomial | log | Total Spore Count | qAIC |
| Pigmented Spore Size | Pigmented Spore Size ~ Geographic Distance + (Geographic Distance)<sup>2</sup> | Gaussian | --- | --- | AIC |
| Pigmented Spore Germination | Germinated Spore Count ~ (Genetic Distance)<sup>2</sup> | Negative Binomial | log | Pigmented Spore Count | qAIC |

**Figure 4.**
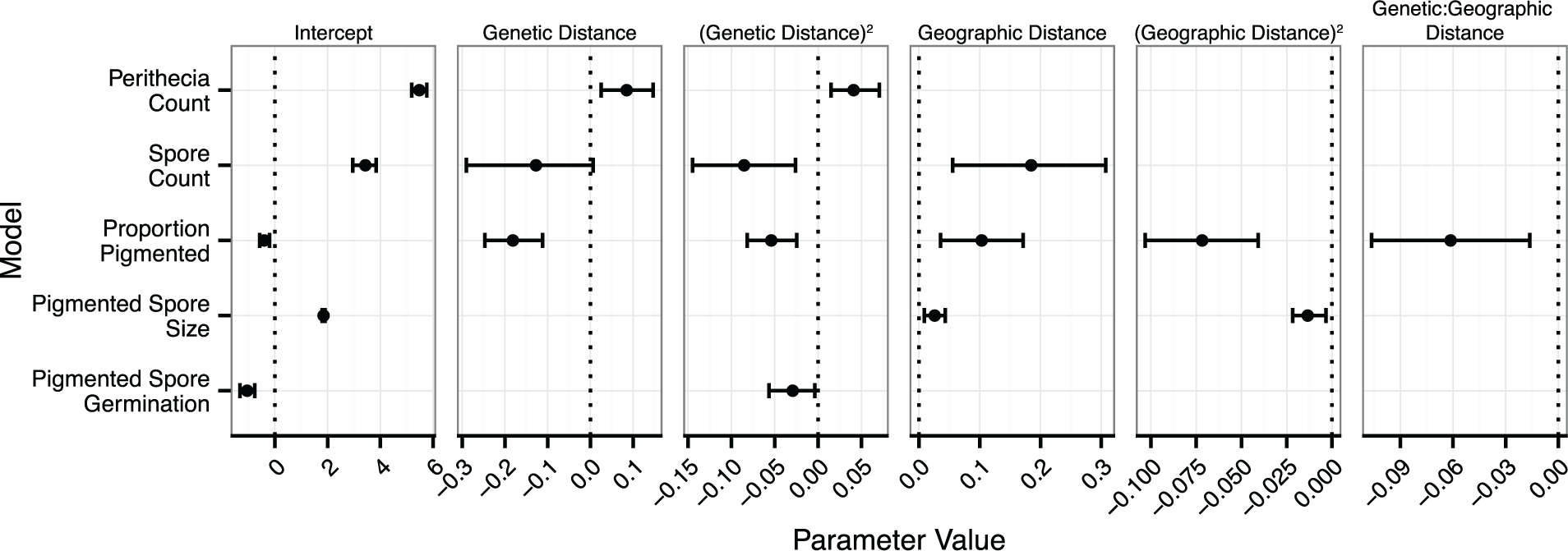
Summary of model parameter values for each measured trait. Parameter values returned from the mixed effects models are represented as points with error bars displaying the 95% Highest Posterior Density (HPD) from 10000 iterations of Markov chain Monte Carlo (MCMC) parameter estimation. Absence of points indicates parameters that were dropped from a model based on qAIC or AIC metrics. Dotted lines indicate the zero values. If the HPD interval overlaps with zero, then the parameter is not different from a null model. Only the genetic distance parameter in the spore count model overlaps with zero. Parameter values for all models except ‘Pigmented Spore Size’ are on a log link scale. Values are listed in Table S1.

The proportions of variance explained by the fixed effects (crossing distance) and the random effects (maternal or paternal parentage), or not explained by the model, vary widely among traits (Fig. 5). However, in every model, the random effects of maternal and paternal strain identity explain more variance than the fixed effects of genetic and geographic distance (Table 4).

**Figure 5.**
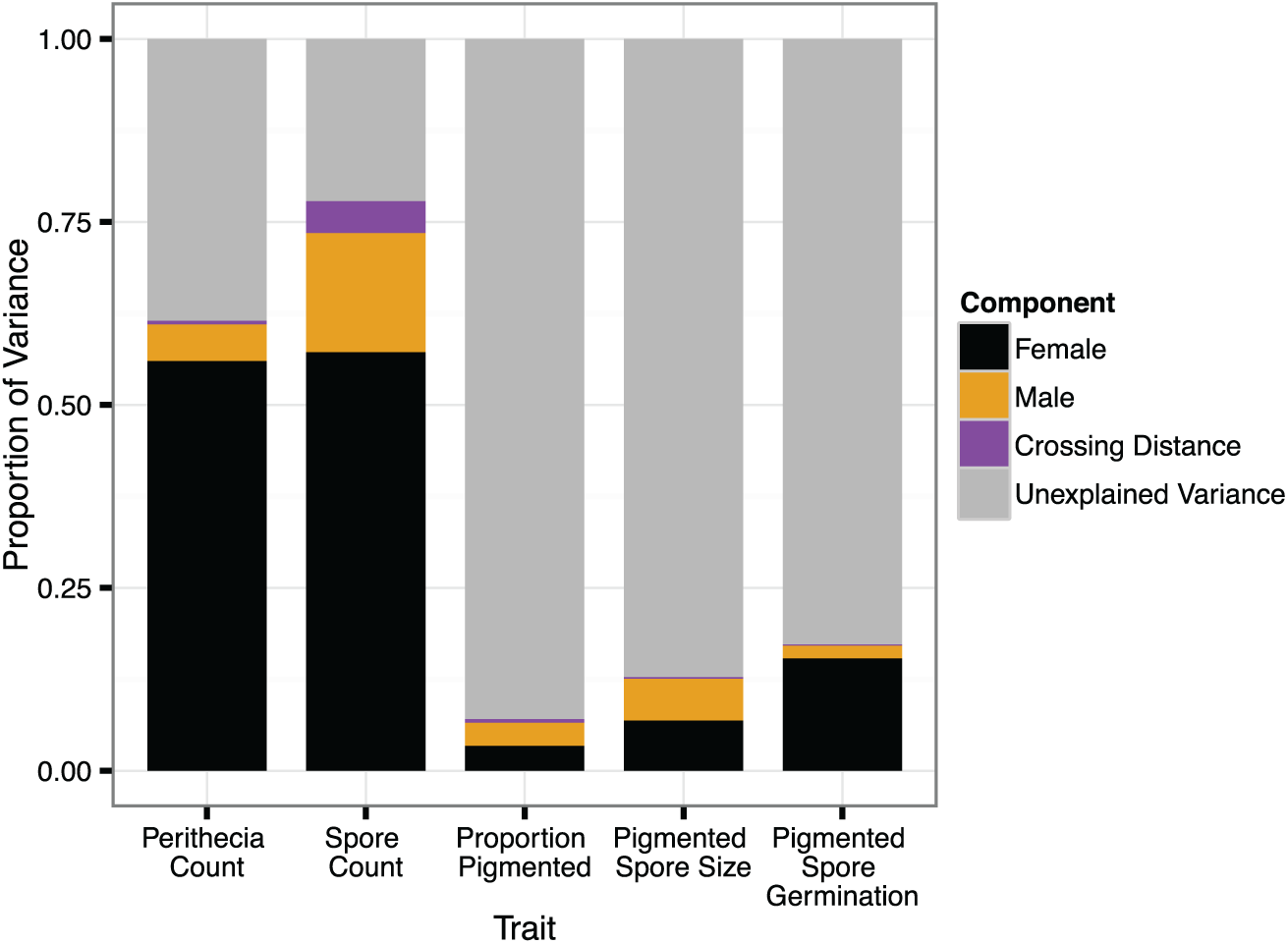
Proportions of variance explained by maternal and paternal random effects, fixed effects (crossing distances), or other unexplained variance. Proportions were calculated using the R^2^GLMM values in Table 4 along with the individual random effect variances returned from the models.

**Table 4.**
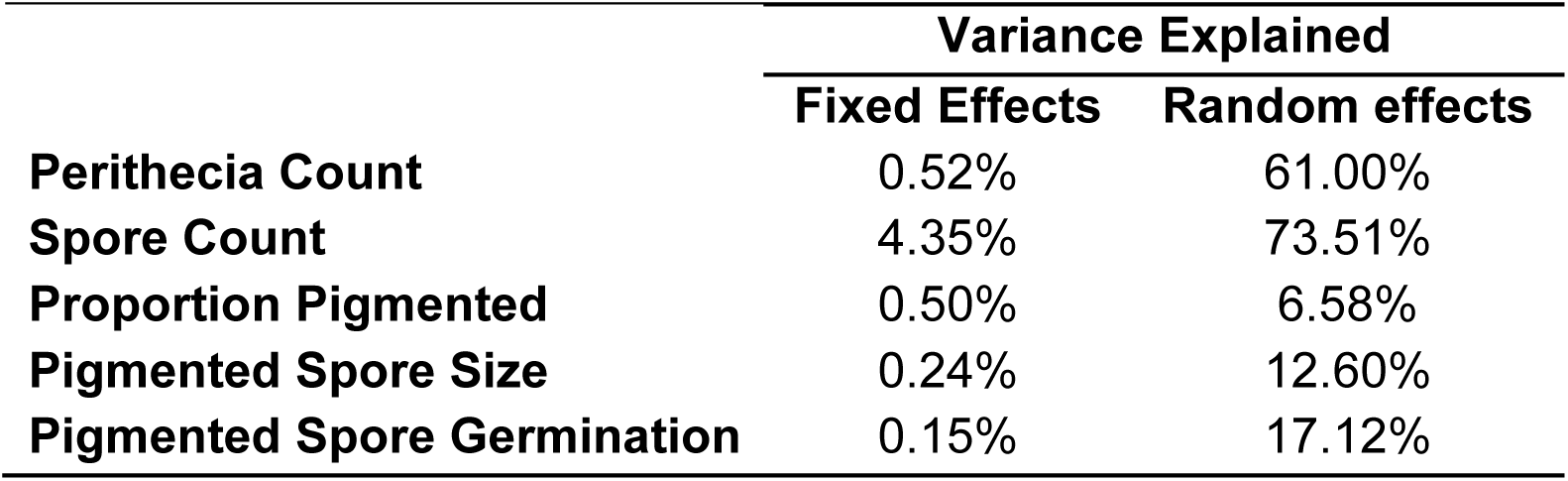
Percentage of variance explained by fixed effects (crossing distances) or maternal and paternal random effects for each modeled trait. Percentages are based on the marginal and conditional R^2^GLMM values calculated for each model (Nakagawa & Schielzeth, 2013).

### Maternal strain identity explains most of the variation in spore production and germination, but not size and pigmentation

Maternal identity explains most of the random effect variation in models of perithecia count, spore count and pigmented spore germination (Fig. 6) and correlations between maternal and paternal BLUPs (the relative influence of a strain as a mother or a father on a given phenotype) for these traits are weak or non-existent (Fig. 7). However, there is no significant difference in the proportion of variance explained by maternal or paternal strains in the models of proportion pigmented and pigmented spore size (Fig. 6) and correlations between maternal and paternal BLUPs for these traits are strong (Fig. 7).

**Figure 6.**
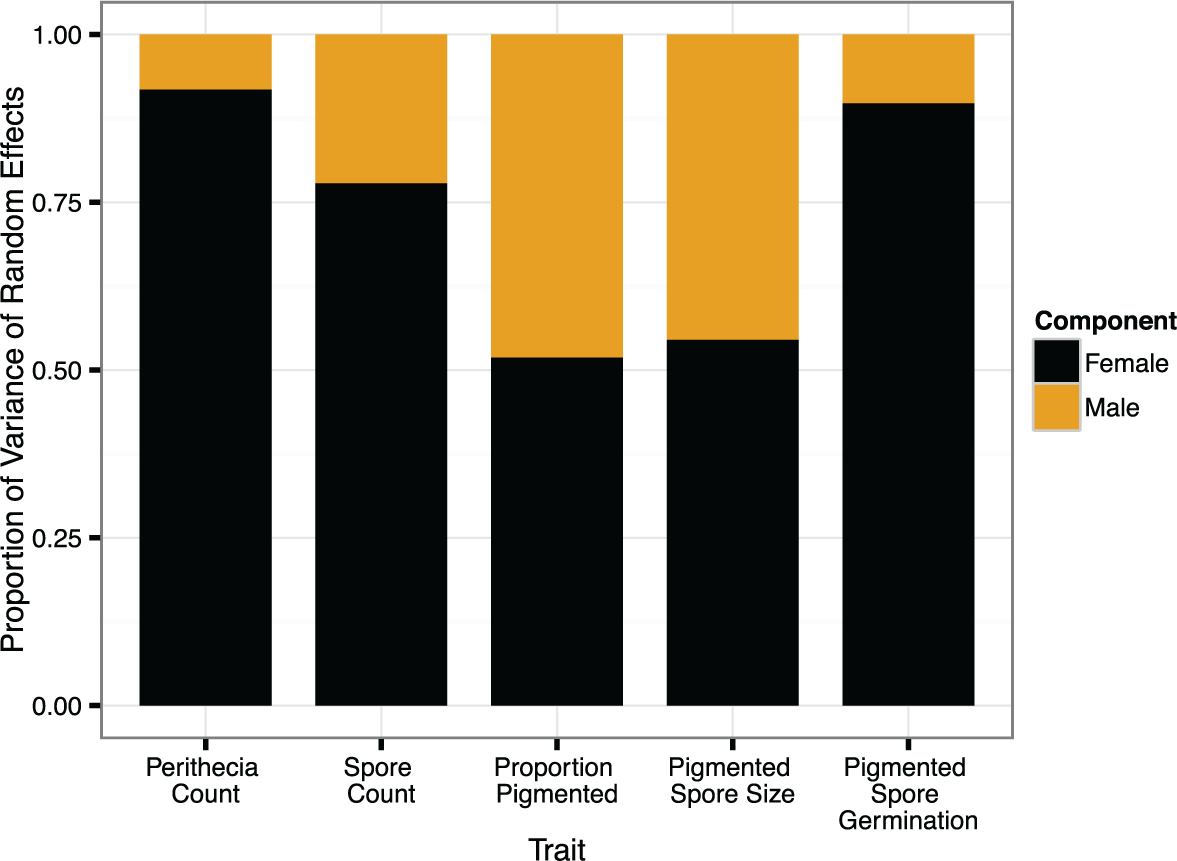
Proportions of random effect variance explained by strains acting as mothers or fathers. Female versus male random effect variances (statistically significant differences in bold): perithecia count ratio of male:female variance = 0.076, *F*_20_,_20_ = 0.07 56, *P* = 3.00e-7; spore count ratio of male:female variance = 0.26, *F*_20_,_20_ = 0.2582, *P* = 0.0039; proportion pigmented ratio of male:female variance = 0.92, *F*_20_,_20_ = .9226, *P* = 0.86; pigmented spore size ratio of male:female variance = 0.82, *F*_20_,_20_ = 0.8206, *P* = 0.66; pigmented spore germination male:female variance = 0.087, *F*_20_,_20_ = 0.08 74, *P* = 1.05e-6.

**Figure 7.**
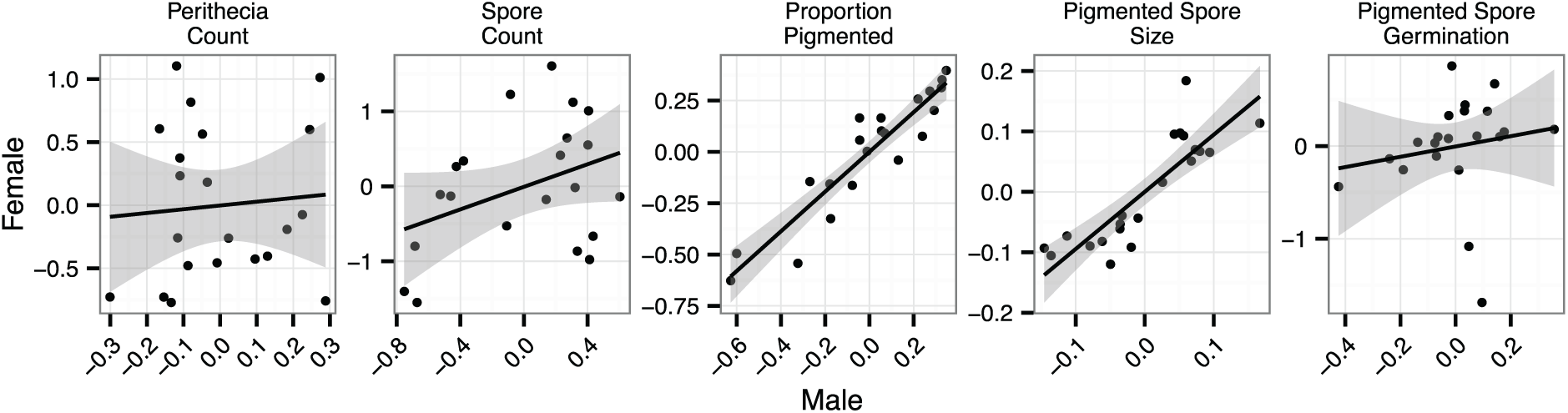
Correlations between male and female BLUPs for all models. Points represent strains. Values are a measure of the relative influence of strains (as mothers or fathers) on phenotype outcomes. Shading shows the 95% confidence interval of the line. Correlations (significant correlations **in bold**): perithecia count r^2^ 0.0067, *F*1,19 = 0.1287, *P* = 0.72; spore count r^2^ = 0.15, *F*_1,19_ = 3.257, *P* = 0.087; proportion pigmented r^2^ = 0.86, *F*_1_,_19_ = 119.5, *P* = 1.23e-09; pigmented spore size r^2^ = 0.73, *F*_1_,_19_ = 52.38, *P* = 7.18e-7; pigmented spore germination r^2^ = 0.027, *F*_1_,_19_ = 0.5259, *P* = 0.48.

## DISCUSSION

Maternal effects emerge as a major feature of the reproductive biology of *Neurospora crassa*. We find strong evidence for maternal effects acting on spore production and spore germination (Fig. 3). While phenotypes are consistent for any particular mother, they vary widely among mothers, and fathers have no consistent influence (Roff, 1998). We find no evidence for strong maternal effects on spore size or the proportion of pigmented spores (Fig. 3).

Our mixed effects models confirm the visual evidence by controlling for potentially important confounding variables: genetic and geographic distances between parents. Crossing distances account for a small proportion of the overall variation in reproductive outcomes, suggesting that inbreeding and outbreeding depression do not substantially influence the success of crosses among our collection of strains (Fig. 5 and Table 4).

In fact, after controlling for the fixed effects of crossing distances in our models, we find that the random effects of parental strain identity account for most of the variance in reproductive outcomes (Fig. 5). Furthermore, variation among strains acting as mothers accounted for most of the explained variation in our models of perithecial production, spore production, and spore germination (Fig. 6). Perithecial production appears to be under strong maternal control (Raju, 1992; Turner *et al.*, 2010), while maternal effects appear to play a dominant role in spore production and spore germination. We distinguish perithecial production as under maternal control—rather than influenced by maternal effects—because maternal effects can only be determined by measuring offspring phenotypes.

Maternal effects do not play a role in spore size or pigmentation; these traits are equally influenced by both parents in a cross, regardless of their roles as mother or father. In fact the effect of each strain acting as a mother is highly correlated with that strain’s effect when it acts as a father (Fig. 7), and mothers exert no more influence than fathers (Fig. 6). Taken together, the analyses suggest that spore size and pigmentation are controlled mainly by direct genetic effects; the large proportion of variance not explained by crossing distance or parental effects for both traits (92.92% and 87.16%) is consistent with direct genetic effects caused by specific allelic variants at one or more loci.

Observed maternal effects are not an artifact of correlations in trait values, and the number of perithicia generated by a cross does not predict subsequent spore phenotypes. Perithecial production, spore production, and spore germination are not correlated with each other (Fig. S4).

### Mapping maternal effects onto reproductive development: A model

The dynamic maternal effects we document are a clear reflection of *N. crassa* ontogeny, and correlate with what is known about the species’ reproduction (Fig. 8). Maternal effects influence spore production, but are not a feature of spore maturation; as spores germinate, maternal effects are observed once again. Spore production takes place within maternal tissues, while spore maturation occurs after spore walls separate developing spores from maternal cytoplasm [38]. Germination draws on resources provided by the mother.

**Figure 8.**
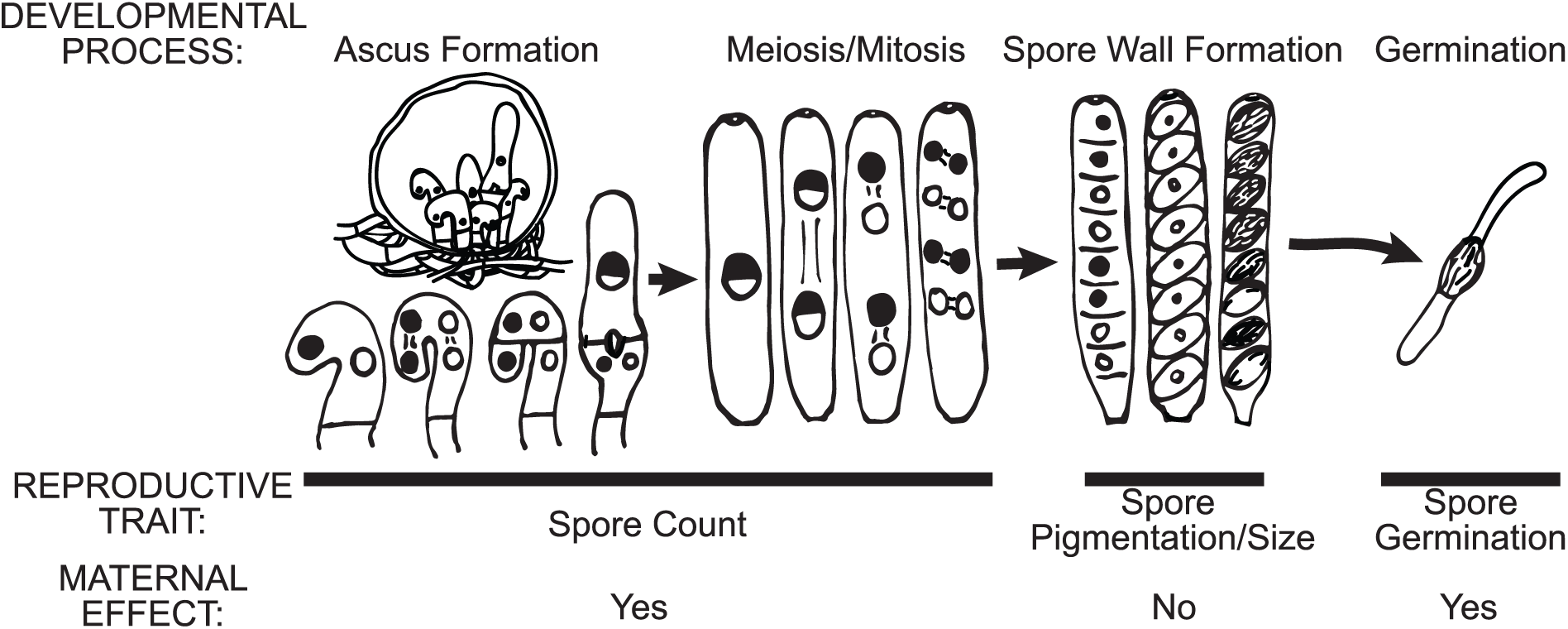
Maternal effects vary dynamically across *N. crassa* reproductive development. Spores are generated within maternally derived asci, and maternal effects play a role in the total number of spores produced. Maternal effects do not influence spore pigmentation or size; these phenotypes develop after spore walls separate the spores from maternal cytoplasm. Maternal effects influence germination; germinating spores use maternally supplied nutrient stores.

The apparent lack of maternal effects associated with spore maturation, and the suggestion that the spore cell wall limits contact between the maternal cytoplasm and the developing spore (Fig. 8), is supported by analyses of spore maturation mutants. These mutants suggest that spore maturation is controlled by the genetic material inherited by each spore (Raju & Leslie, 1992). Genes appear to be expressed autonomously within each of the spores of an ascus (Freitag *et al.*, 2004). Both studies suggest mothers have limited control over spore maturation, instead, the genetic material of the spores themselves appears to be in direct control of pigmentation and size.

Mechanisms causing the maternal effects associated with the germination of pigmented spores are less clear, but possible conduits include the carbohydrates and lipids packaged within spores. The raw materials used to form a spore are derived from the maternal parent. Germinating spores use stored materials for energy production during germination, and the amount of lipids in a spore can influence germination (Lingappa & Sussman, 1959; Budd *et al.*, 1966). Maternal strains may differ in their ability to supply carbohydrates and lipids to developing spores, resulting in maternal effects. Maternal effects may also be mediated through mitochondria. Mitochondrial influences do not qualify as maternal effects under at least one definition (Wolf & Wade, 2009), because mitochondrial DNA is a property of the offspring and not the mother. However, in germinating spores of *Neurospora tetrasperma,* a species closely related to *N. crassa* (Dettman *et al.*, 2003a), the maternal mitochondria are responsible for all energy synthesis until 90 mins. after germination initiates, at which point a germinating spore begins to synthesize its own mitochondria (Hill *et al.*, 1992). Germination success may be influenced by both the maternal provision of nutrients, as well as maternal supply of mitochondria.

### Potential for maternal effects in other fungi

Maternal effects are likely to influence the life histories of other anisogamous ascomycetes (Billiard *et al.*, 2010), as well as anisogamous chytrids, including species of *Allomyces* and *Monoblepharis* (James *et al.*, 2006). Maternal effects may also play a role in basidiomycete life histories. While the majority of basidiomycetes do not produce specialized sexual structures, when small and large haploid mycelia fuse, disproportionate cytoplasmic inheritance (from the larger mycelium) can result (Griffiths, 1996).

### Anisogamy and maternal effects in diverse eukaryotes

Anisogamy is commonly referenced as the basis for maternal effects and is widespread among eukaryotes, but maternal effects are thoroughly studied in only two kingdoms—animals and plants. Our discovery of maternal effects in *N. crassa,* an anisogamous fungus, suggests anisogamy as a strong predictor of maternal effects across the eukaryotic domain.

Other anisogamous species are found in groups including apicocomplexans, dinoflagellates, diatoms, parabasalans, and red algae (Dacks & Kasinsky, 1999). The red algae in particular have life cycles very similar to filamentous anisogamous ascomycetes (Demoulin, 1985). Cytoplasmic inheritance in the malaria parasite *Plasmodium falciparum,* an anisogamous apicocomplexan, is predominantly female (Vaidya *et al.*, 1993). We hypothesize that maternal effects occur in, and have important implications for the evolution of, all of these anisogamous taxa.

### Conclusions

A simple crossing experiment suggests maternal effects profoundly influence fundamental aspects of individual fitness in the genetic model *N. crassa.* The maternal influence on spore production and viability is stronger than the influence of genetic divergence or geographic isolation—two essential aspects of any cross thought to influence reproductive success and speciation in all sexually reproducing organisms. Our experiment increases the number of eukaryotic clades known to house organisms influenced by maternal effects from two, the plants and animals, to three—the plants, animals, and fungi. Given the distribution of anisogamy across many other eukaryotic clades, and the correlation between anisogamy and maternal effects, we hypothesize that maternal effects influence the evolution and ecology of additional, as yet untested, eukaryotic organisms.

## ACKNOWLEDGEMENTS

We thank J. Paltseva for assistance in counting perithecia, P. Rogers for assistance with flow cytometry, R. Corbett-Detig for help with genetic distance calculations, J. Taylor for clarification of SNP data, and M. Lalli for critical reading of this manuscript. We are also grateful for the resources provided by the Fungal Genetics Stock Center (Manhattan, Kansas, USA). Our work is supported by the National Science Foundation Graduate Research Fellowship awarded to K.Z. and by other National Science Foundation grants awarded to the Pringle Laboratory. This work was also supported by funds from the Max Planck Institute for Demographic Research to K.Z., D.L., and A.P. The work of D.L. was supported by the Max-Planck Odense Center, a collaboration of the Max Planck Society and the University of Southern Denmark.

## SUPPLEMENTARY INFORMATION

**Table S1. Summary of model parameter values for each measured trait.**

**Figure S1. Example plot of forward scatter height (FSC-H) vs. side scatter width (SSC-W) of clustered flow cytometry data from a pooled set of 96 analyzed ascospore samples.**

**Figure S2. Asymptotic non-linear least squares regression of the proportion of pigmented spores measured by flow cytometry *vs*. counting.**

**Figure S3. Asymptotic non-linear least squares regression of the proportion of germinated spores measured by counting *vs*. flow cytometry. !**

**Figure S4. Correlogram showing all possible pairwise comparisons between the five measured traits.**

**Analysis S1. PDF file showing R code and output for all analyses and figures in main text.**

**Derivation S1. Derivation of distribution specific variance for the negative binomial distribution.**

